# Alterations in sperm long RNA contribute to the epigenetic inheritance of the effects of postnatal trauma

**DOI:** 10.1101/386037

**Authors:** K. Gapp, G. van Steenwyk, P.L. Germain, W. Matsushima, K.L.M. Rudolph, F. Manuella, M. Roszkowski, G. Vernaz, T. Ghosh, P. Pelczar, I.M. Mansuy, E.A. Miska

**Author notes:** These authors share last authorship (I.M. Mansuy, E.A. Miska). Correspondence to IMM or EAM.

## Abstract

Psychiatric diseases have a strong heritable component known to not be restricted to DNA sequence-based genetic inheritance alone but to also involve epigenetic factors in germ cells ^1,2^. Initial evidence suggested that sperm RNA is causally linked ^2,3^ to the transmission of symptoms induced by traumatic experiences. Here we show that alterations in long RNA in sperm contribute to the inheritance of specific trauma symptoms. Injection of long RNA fraction from sperm of males exposed to postnatal trauma recapitulates the effects on food intake, glucose response to insulin and risk-taking in adulthood whereas the small RNA fraction alters body weight and behavioral despair. Alterations in long RNA are maintained after fertilization, suggesting a direct link between sperm and embryo RNA.

## Introduction

Adverse experiences can have long-lasting transgenerational effects on mental and physical health, and often increase disease risk ^4,5^. Traumatic stress in early life in particular, can induce pathologies like psychosis, depression and metabolic dysfunctions in adulthood across generations ^6^. To examine the biological factors involved, we recapitulated heritable behavioural and metabolic effects of postnatal trauma across several generations using a previously established model of unpredictable maternal separation combined with unpredictable maternal stress (MSUS) in the mouse, that shows symptoms through up to three generations (Fig. 1) ^2,7–13^. We have shown that such postnatal trauma alters small RNA in sperm and that injection of total sperm RNA from exposed male mice into naïve fertilized oocytes elicits symptoms reminiscent of those observed in natural offspring of exposed fathers ^2^. Other studies have demonstrated that adult stress ^14,15^ and environmental insults like altered diet or vinclozolin exposure, or positive factors such as exercise or environmental enrichment can affect small RNA in sperm ^16–21^ and somatic tissues ^22^ in the offspring. Recently, tRNA fragments and their modifications were also found to be affected by nutritional insult, and unmodified or modified sperm small RNA injected into fertilized oocytes could mimic metabolic changes resulting from altered parental diet in the progeny ^18,23,24^. These studies therefore suggest that small RNA in sperm can be carrier of heritable information. Here we sought to determine whether long RNA in sperm also contributes to the transmission of the effects of previous exposure.

**Figure 1.**
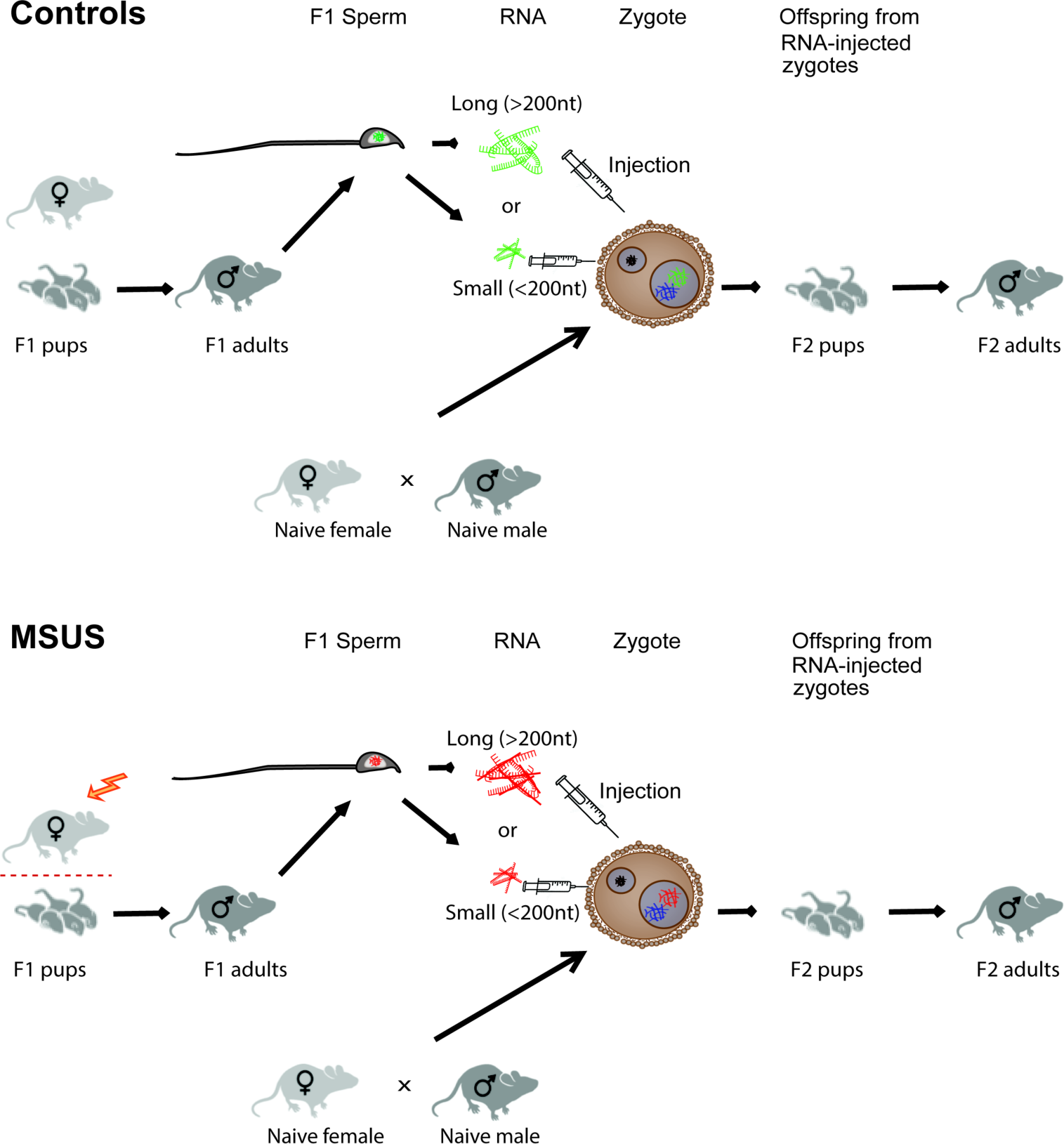
Generation of control and MSUS F1 males and of animals resulting from sperm RNA injections into zygotes. Between postnatal day (PND) 1 to 14, mouse pups are exposed to unpredictable maternal separation combined with unpredictable maternal stress (MSUS) while control mice are left undisturbed. When adult, sperm of F1 MSUS and control males is harvested, RNA is extracted, size selected then used for injection into naïve fertilized oocytes. Long RNA (>200 nt) includes mRNA, lncRNA, TE RNA among others. Small RNA (<200 nt) includes miRNAs, tRNA fragments, piRNAs among others. Injected zygotes are implanted into foster mothers to produce offspring. (nt = nucleotides).

## Materials and Methods

### Mice

C57BL/6J mice were housed in a temperature and humidity-controlled facility under a reverse light-dark cycle, and food and water were provided *ad libitum*. Experimental procedures were performed during the animals’ active cycle. All experiments were approved by the cantonal veterinary office, Zurich (license 55/12 then 57/15).

### MSUS paradigm

C57BL/6J primiparous females and males were mated at 2.5-3 months of age. Randomly selected dams and litters were subjected to daily 3h proximal unpredictable separation combined with unpredictable maternal stress (MSUS) from postnatal day 1 to 14 as described previously ^13^ or left undisturbed (Controls). Cage changes took place once a week until weaning (postnatal day 21) for both MSUS and control animals. At weaning, pups were assigned to social groups (4-5 mice per cage) of the same treatment and sex, but from different dams to prevent litter effects.

### Sperm collection

Males used for sperm collection did not undergo any behavioural or metabolic testing and were approximately 3 months old at the time of sampling. For sperm RNA sequencing, mature sperm cells were collected from cauda epididymis and purified by counterflow centrifugal elutriation using a Beckman JE-5.0 elutriation rotor in a Sanderson chamber and a Beckman Avanti J-26 XPI elutriation centrifuge as described previously ^2^. Purity of samples was confirmed by inspecting eluted sperm cells under a light microscope. For RNA injection experiments, sperm cells were separated from potential somatic contamination by swim-up ^39^.

### Zygotes collection

C57Bl/6J primiparous females underwent superovulation consisting of intraperitoneal injection of 5 IU pregnant mare serum gonadotropin followed by 5 IU human chorionic gonadotrophin 48 h later. Females were then left to mate with either an age-matched MSUS or control male. After plug detection, zygotes were scraped out of uterine horn and collected directly into Trizol LS solution.

### RNA extraction

Total RNA was prepared from adult mouse sperm using a standard Trizol protocol. Total RNA was prepared from zygotes and 4-cell embryos using the Trizol LS protocol. The quantity and quality of RNA were determined by Agilent 2100 Bioanalyser (Agilent Technologies) and Qubit fluorometer (Life Technologies).

### RNA size separation

500 ng sperm RNA pooled from 4 control or 4 MSUS males each were subjected to Agencourt AMPure XP (Beckman Coulter) following the manufacturer’s instructions. Total RNA contained in 50 ul water was mixed with 125ul AMPure XP bead solution, incubated for 5 minutes at room temperature then placed on a magnetic stand to allow magnetic beads to bind long RNA. When the solution appeared clear, the supernatant containing small RNA was aspirated and saved for later processing. Beads with bound long RNA were washed twice with 200ul of 70% EtOH. Long RNA were eluted by addition of 100ul elution buffer. Samples were placed again on the magnetic stand to capture magnetic beads while aspirating the supernatant containing long RNA and transferring it to a new tube for subsequent processing. Small and long RNA fractions were subjected to EtOH precipitation for purification. Efficiency of size separation was determined using Agilent 2100 Bioanalyzer (Supplementary Figure 2a).

### RNA fragmentation

130 ul of long RNA in Tris-HCL, pH 8.0 buffer was transferred into Crimp cap Micro tubes and subsequently fragmented using a Covaris Ultrasonicator E210 containing an AFA intensifier using the following settings: time: 450sec, 175 Watt, 200 cycles/burst, intensity 5, duty cycle 10%, temperature 4-8°C, Power mode: frequency sweeping, degassing mode: continuous, water lever: 6. Successful fragmentation was determined using Agilent 2100 Bioanalyzer (Supplementary Figure 2b).

### RNA injection in fertilized oocytes and embryo culture

Control fertilized oocytes were collected from C57BL/6J females (Janvier, France) after superovulation (intraperitoneal injection of 5 IU pregnant mare serum gonadotropin followed by 5 IU human chorionic gonadotrophin 48h later), and mating with C57BL/6J males. 1-2 pl of 0.5 ng/µl solution of small or long RNA isolated and pooled from sperm from of 4 adult MSUS or control males dissolved in 0.5 mM Tris-HCl, pH 8.0, 5 µM EDTA were microinjected into the male pronucleus of fertilized oocytes using a standard microscope and DNA microinjection method ^40^. Injection of fragmented long RNA from sperm used the same protocol and the same samples but fragmentation. This amount of injected RNA corresponds to the amount of sperm RNA estimated to be delivered to the oocyte by a sperm cell ^41^. The experimenter was blind to treatment (double blinding method), and injections were balanced across groups and were all performed between 2 and 4 pm. For molecular analysis at the 4-cell stage, embryos were placed in KSOM medium (EmbryoMax Powdered Media Kit, Millipore Cat# MR-020P-5F) supplemented with essential amino acids (Millipore) and cultured for 48h in 5% CO2 at 37C°.

### Behavioural testing

The experimenter was double-blinded to treatment, meaning the group assignment was coded twice prior to experiments, once by the experimenter preparing RNA samples and once by the experimenter performing RNA injections. Behaviours were videotaped and scored both manually and automatically by tracking software (Viewpoint). All behavioural tests were conducted in adult age-matched male animals.

#### Elevated plus maze

The elevated plus maze consisted of a platform with two open arms (no walls) and two closed arms (with walls) (dark gray PVC, 30×5 cm) elevated 60 cm above the floor. Mice were placed in the center of the platform facing a closed arm, and tracked for 5 minutes. The latency to enter an open arm was manually scored and total distance moved was automatically recorded by a videotracking system.

#### Light-dark box

Each mouse was placed in the lit compartment (white walls, 130 lux) of a box (40×42×26 cm) split into two unequal compartments (2/3 lit, 1/3 dark compartment with black walls and covered by a black lid and by a divider with a central opening (5×5 cm). Mice could freely move from the lit to the dark compartment during a 10-min period. Time spent in each compartment and latency to enter the dark compartment were measured manually.

#### Forced swim test

Mice were placed in a beaker of cold water (18+/−1°C) for 6 min. Floating duration was scored manually.

### Metabolic testing

Experimenters were double-blinded to treatment, meaning both the experimenter conducting RNA injections and the experimenter conducting phenotyping on resulting offspring was not aware of groups assignment. All metabolic tests were conducted in adult age-matched males.

#### Caloric intake measurement

The amount of consumed food was measured for each cage in 4 months old animals every 24h. Caloric intake was calculated as the mean amount of food intake over 72h in relation to mean body weight (caloric intake = mean food intake/mean body weight).

#### Glucose (GTT) and insulin (ITT) tolerance test

Mice were fasted for 5h. For GTT, glucose was measured in blood at baseline and 0, 15, 30 and 90 min after intraperitoneal injection of 2 mg per g body weight of glucose in sterile 0.45% (wt/vol) saline (injection started at 2 pm). For ITT, glucose was measured in blood at baseline, and 0, 15, 30, 90 and 120 min after intraperitoneal injection of 1 mU per g body weight of insulin (NovoRapid Novo Nordisk A/S) in sterile 0.9% saline (injection started at 2 pm). If blood glucose decreased below 1.7 mM/ml, it was rescued by intraperitoneal injection of 2 mg/g of glucose. For both GTT and ITT, glucose level was determined in fresh tail blood using an Accu-Chek Aviva device (Roche).

### RNA sequencing (RNAseq)

Sequencing was done using an Illumina Genome Analyzer HiSeq 2500 (Illumina) in Rapid run mode for 51 cycles plus 7 cycles to read the indices in two separate runs (run 1 consisted of libraries representing biological replicates of control sperm RNA each pooled from 5 males and libraries representing biological replicates of sperm RNA from males exposed to MSUS, each pooled from 5 males. Run 2 consisted of technical replicates of run 1 plus one additional sample representing sperm RNA from 5 males exposed to MSUS. RNA libraries for first sequencing run were prepared with the NEBNext Ultra Directional RNA Library Prep Kit (NEB#E7420S, New England BioLabs Inc.) following the manufacturer's recommendations. Libraries for the second run were prepared using the TruSeq Stranded Total RNA kit according to the manufacturer's instructions with indices diluted at 1:3. 220 ng of total sperm RNA was subjected to removal of rRNA using Ribozero (first run) and Ribozero gold kit (second run). Efficiency of Ribozero treatment was validated by Agilent 2100 Bioanalyzer. An average of 6.4 ng of zygotic RNA and 3.7 ng of 4-cell embryo RNA per sample were not subjected to ribosomal depletion but immediately used for first strand synthesis. After demultiplexing and adaptor removal, an average of 60,948,122 pass filter reads was obtained in each sperm library and 43,346362,5 in the zygotes libraries.

### Bioinformatic analysis

RNAseq reads were trimmed of adapter sequences on the 3’ end using cutadapt v1.14 with a 5% error rare and using adapter sequences of the Truseq universal adapter, the Truseq indexed adapters (with sample-specific barcodes) and their reverse complement. Reads with a length of less than 15bp were discarded. Samples were then directly quantified using Salmon v0.9.1 ^42^ on an index created from the GRCm38 genome using, as transcripts: 1) GENCODE vM16 features, 2) transposable elements from repeat masker (concatenating stranded features associated to the same family of elements), and 3) piRNA precursors ^43^. Given the weak strand bias (roughly 65/35) of the libraries, they were treated as unstranded. RNA composition estimates (Supplementary Figure 4) were obtained by summing the reads of transcripts belonging to the same RNA biotype and dividing by the total number of assigned reads in the sample. For all downstream analyses, transcript counts were then summed by gene symbol. Expression profiles during spermatogenesis and from Sertoli cells were obtained from the Gene Expression Omnibus (respectively GSE100964 and GSM1069639) ^28,29^ and quantified as indicated above. Library type was specified as respectively SF (strand-specific reads coming from the forward strand) and ISR (inward strand-specific reads coming from the reverse strand). For differential expression analysis, only lincRNAs and protein-coding genes that had at least 20 reads in at least 3 samples were tested. Differential expression analysis was performed with edgeR, using the exact test for zygotes and 4-cell embryos, while for sperm, generalized linear models were used to account for the technical differences between batches (~batch+condition). Given the presence of technical replicates in the sperm dataset, we performed two additional control analyses showing that largely the same results could be obtained by either, i) using only the most recent batch (Supplementary Figure 7a), or ii) accounting for the incomplete independence of the samples as described in ^44^ (Supplementary Figure 7b). Gene ontology enrichment analysis was performed using the goseq R package ^45^ (Fisher’s exact test) to account for the length bias in RNAseq experiments, using GO terms with 10 to 1000 annotated genes. Only genes with existing GO annotations were used, and for enrichments analysis performed on differentially expressed genes, only tested genes (i.e. after the aforementioned count filtering) were used as background. When indicated, most specific enrichments were obtained by removing from the results, terms that had significantly enriched related sub-terms. For heatmaps and PCA, log(1+normalized counts) were used. Small RNAseq libraries were pre-processed with cutadapt to remove adapters. Subsequently, only reads ending in CCA-3ʹ were selected for enrichment analysis of tRNA-derived fragments. CCA-3ʹ was trimmed off, and reads were quantified using Salmon. For this, an index of tRNA sequences was constructed using the mouse tRNA sequences from GtRNAdb ^46^ with parameters “--perfectHash --kmerLen 15”. Differential expression analysis was performed using DESeq2 and tximport ^47,48^. tRNA gene loci were grouped into genes by trimming off the trailing number from the GtRNAdb annotation names, and tRNA with pseudo-counts equal to 0 in all replicates were removed from the analysis prior to running DESeq2 with default parameters.

### Statistical analyses

Samples size was estimated based on previous work on the MSUS model ^2,9–13^. Two-tailed Student *t* tests were used to assess statistical significance for behavioural, body weight, caloric intake and GTT measurements. Chi-square test for 2 binominial populations with Yates continuity correction was used to analyze the proportion of animals rescued on the ITT before 90 minutes post-injection. When data did not match the requirements for parametric statistical tests (normal distribution), Mann Whitney U test was applied. If variance was not homogenous between groups, adjusted *P* value, *t* value and degree of freedom (Welch correction) were determined in parametric tests. Data were screened for outliers using prism`s ROUT test (for total distance moved on the elevated plus maze, time spent in bright on light-dark box, time spent floating on forced swim test, weight, food intake and glucose response) and identified animals were excluded from analysis. All statistics of behavioural and metabolic tests were computed with Prism and SPSS. All reported replicates were biological replicates, or pooled samples from biological replicates in the case of sequencing samples and RNA injection samples. Significance was set at *p* < 0.05 for all tests. Boxplot whiskers represents the Tukey method.

### Data availability

Small RNAseq data have been deposited in the ArrayExpress database at EMBL-EBI (www.ebi.ac.uk/arrayexpress) under accession number E-MTAB-5834 (sperm), E-MTAB-6589 (zygotes) and E-MTAB-6587 (4-cell embryos). All relevant data are available from the authors. Previously published datasets used for comparisons in this study are available under Gene Expression Omnibus (respectively GSM1069639, GSE100964 and GSE50132) ^2,28,29^.

### Code availability

Code used to generate results is available from the authors upon request.

## Results

The impact of MSUS on behaviour. First, using the MSUS paradigm, we produced a cohort of mice exposed to traumatic stress in early postnatal life and confirmed their behavioural phenotype in adulthood described previously ^2,13^ (Supplementary Figure 1). As expected, F1 MSUS adult males had increased risk-taking behaviours reflected by shorter latency to first enter an open arm on an elevated plus maze, and more time spent in the bright field of a light dark box (Figs.S1a,c) with increased total locomotor activity (Supplementary Figure 1b) but no change in latency to first enter the dark compartment of the light dark box (Supplementary Figure 1d). They also had a tendency for increased behavioural despair shown by more time spent floating on a forced swim test (Supplementary Figure 1e).

### The effects of small versus long RNA from sperm on behaviour and metabolism

We then harvested sperm from adult control and MSUS males, extracted total RNA and fractionated the RNA into small (<200 nucleotides (nt)) and long (>200 nt) RNA using Ampure beads. Effective separation was confirmed by Bioanalyzer analysis (Supplementary Figure 2a). We then used these fractions for pronuclear injection into fertilized oocytes. Injection of the small or long RNA fraction from MSUS sperm replicated different hallmarks of the MSUS phenotype in the resulting offspring ^13^ (Figs. 2,3, S3a,b). Injection of small or long RNA alone was insufficient to reproduce changes in risk-taking behaviors on the elevated plus maze observed in natural MSUS animals (Fig. 2a) with small RNA inducing a decrease in locomotor activity inconsistent with natural MSUS offspring ^2^. However, injection of long but not small RNA induced a tendency for increased time spent in the bright field of the light dark box (Fig. 2b), similar to natural MSUS offspring, whereas the injection of small RNA resulted in a tendency for a decreased time spent in the bright field. The small RNA fraction mimicked increased behavioural despair on the forced swim test (Fig. 2c) ^2^. Thus, analysis of the behavioural phenotype in the offspring resulting from sperm RNA injections into zygotes indicate the necessity of both RNA fractions to mimic behavioural changes known to be induced by MSUS in natural offspring.

**Figure 2.**
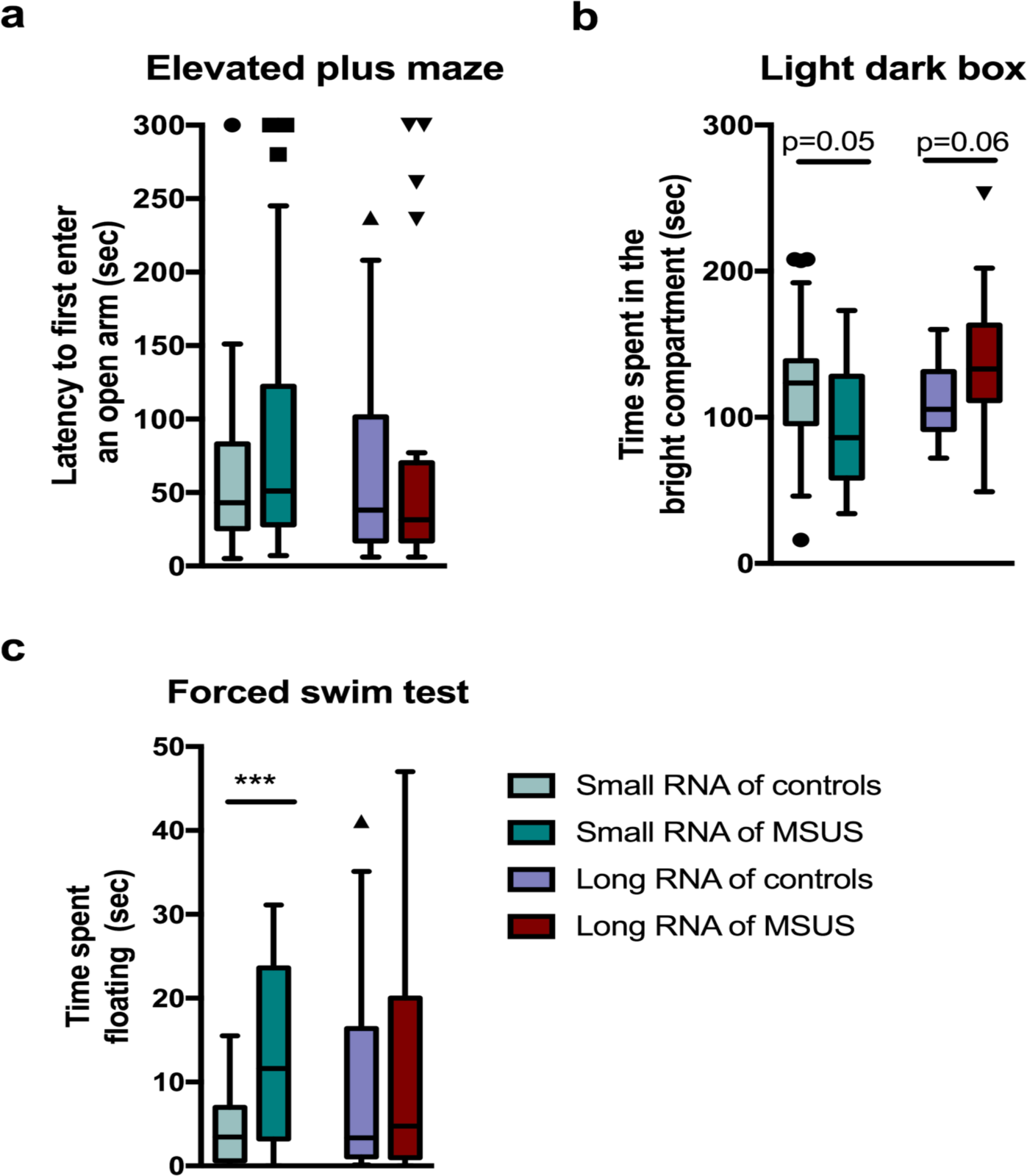
The long RNA fraction from sperm of males exposed to MSUS is sufficient to mimic some behavioural alterations observed in natural offspring of MSUS fathers. Injection of small or long sperm RNA of F1 MSUS males into naïve zygotes has no effect on (a) the latency to first enter an open arm on an elevated plus maze (Small RNA of controls n=25, Small RNA of MSUS n=20, Mann Whitney U=214.5, p>0.05); Long RNA of controls n=17, Long RNA of MSUS n=22, Mann Whitney U=171, p>0.05). Injection of small sperm RNA but not long sperm RNA increases (b) time spent in the bright compartment in a light dark box (Small RNA of controls n=24, Small RNA of MSUS n=17, t(39)=2,02 p=0.05; Long RNA of controls n=18, Long RNA of MSUS n=21, t(32,19)=-1.95, p>0.05) and (c) time spent floating minutes 3 to 6 on a forced swim test (Small RNA of controls n=20, Small RNA of MSUS n=19, t(22.59)=-3.84, p=0.001; Long RNA of controls n=16, Long RNA of MSUS n=22, Mann Whitney U=158, p>0.05) in the resulting male offspring in adulthood. Data are median ± whiskers. Dots, boxes and triangles: values that lie outside the sum of the 75^th^ percentile and 1.5 x the interquartile range or the 25^th^ percentile minus 1.5 x the interquartile range (all values included in statistical analysis). *p<0.05, ***p<0.001.

Further, injection of small RNA increased body weight (Fig. 3a) contrary to that in natural MSUS offspring, which has lower body weight. Long RNA increased food consumption (Fig. 3b) and the sensitivity to insulin challenge (Fig. 3c), similar to natural MSUS offspring ^2^. Injection of small or long RNA fraction alone was insufficient to elicit altered glucose clearance in response to a glucose challenge (Fig. 3d), indicating again, that both small and long RNA together are required to mimic all aspects of the metabolic changes. Given the fact that total RNA injection does mimic alterations observed in natural MSUS offspring ^2^, these results strongly suggest that alterations in both small RNA and long RNA together mediate the effects of postnatal trauma from father to offspring, while alone, they do not or can even cause different effects.

**Figure 3.**
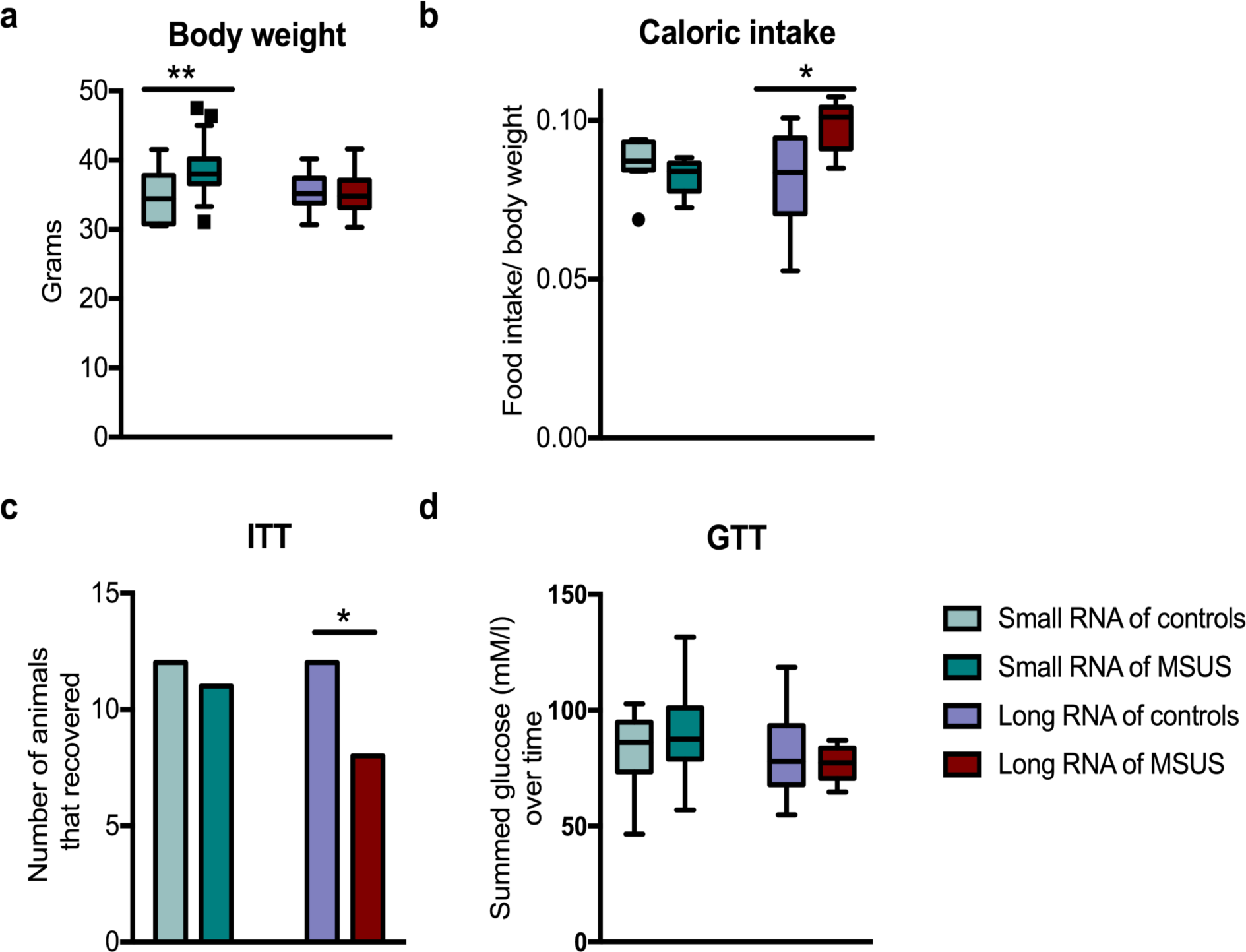
The long RNA fraction from sperm of males exposed to MSUS is sufficient to mimic some metabolic alterations observed in natural offspring of MSUS fathers. Injection of sperm small RNA of F1 MSUS males into naïve zygotes (a) decreases body weight (Small RNA of controls n=16, Small RNA of MSUS n=19, t(33)=3, **p<0.01; Long RNA of controls n=15, Long RNA of MSUS n=21, t(34)=0.37; p>.05). Injection of long RNA (b) increases average caloric intake measured over 72 hours (Small RNA of controls n=8, Small RNA of MSUS n=6, t(12)=1.08, p>0.05; Long RNA of controls n=8, Long RNA of MSUS n=6, t(12)=2.29; p<0.05) and (c) increases sensitivity to insulin after an insulin challenge (Small RNA of controls n=12, Small RNA of MSUS n=12, *X^2^* (1, n=24)=0.6, p>0.05; Long RNA of controls n=12, Long RNA of MSUS n=12, *X^2^* (1, n=24)=2.7, p=0.05). (d) Neither RNA fraction affects the glucose response after a glucose challenge (Small RNA of controls n=12, Small RNA of MSUS n=12, t(22)=1.17; Long RNA of controls n=12, Long RNA of MSUS n=12, t(22)=0.77; p>0.05) in the resulting male offspring when adult. Data are median ± whiskers. Triangles: values that lie outside the 25^th^ percentile minus 1.5 x the interquartile range (all values included in statistical analysis). *p ≤0.05, **p<0.01.

### Consequences of MSUS on sperm transcriptome

Assessment of the sperm transcriptome by next-generation sequencing revealed that long RNA accumulated in mature sperm is dramatically altered in adult males exposed to MSUS compared to control conditions. Besides ribosomal RNA, mitochondrial ribosomal RNA and repeat elements reads mapping to coding and non-coding regions could be detected, consistent with previous reports ^25^ (Figs.S4). The transcripts giving rise to the detected reads were intact and not fragmented as indicated by their distribution spanning the entire transcripts size range (Supplementary Figure 5) and the expected RNA size profile observed by bioanalyzer (Supplementary Figure 2a). Gene ontology (GO) term enrichment analysis of reads mapping to genes in control sperm revealed enrichment for RNA processing, cellular macromolecular complex assembly and chromatin organization among others (Table S1), suggesting gene and transcript regulatory functions. Principal component analysis (PCA) revealed that sperm samples from control and MSUS males segregate (Supplementary Figure 6).

Further quantitative comparison of long RNA from control and MSUS sperm showed significant differential load of several mRNAs and long intergenic non-coding RNAs (lincRNAs) (Fig. 4a,b). GO term analysis revealed an enrichment for cell adhesion and extracellular matrix organization, among others (Fig. 4c, Table S2, Supplementary Figure 7). Interestingly, reads mapping to transposable elements (TEs) were dysregulated in the sperm of MSUS males (Supplementary Figure 8a). When analyzing TEs separately, we found that the relative abundance of several relatively young retro-TEs ^26^ is higher in sperm of males exposed to MSUS (Supplementary Figure 8b). Whether this indeed reflects an activation of these TEs in response to MSUS remains to be determined. Consistent with the induction of some aspects of the MSUS phenotype by sperm small RNA (Fig. 3c), we previously demonstrated sperm transcriptomic alterations in miRNA and piRNA ^2^. Surprisingly, tRNA 3’ fragments were not significantly altered in MSUS sperm (Supplementary Figure 9).

**Figure 4.**
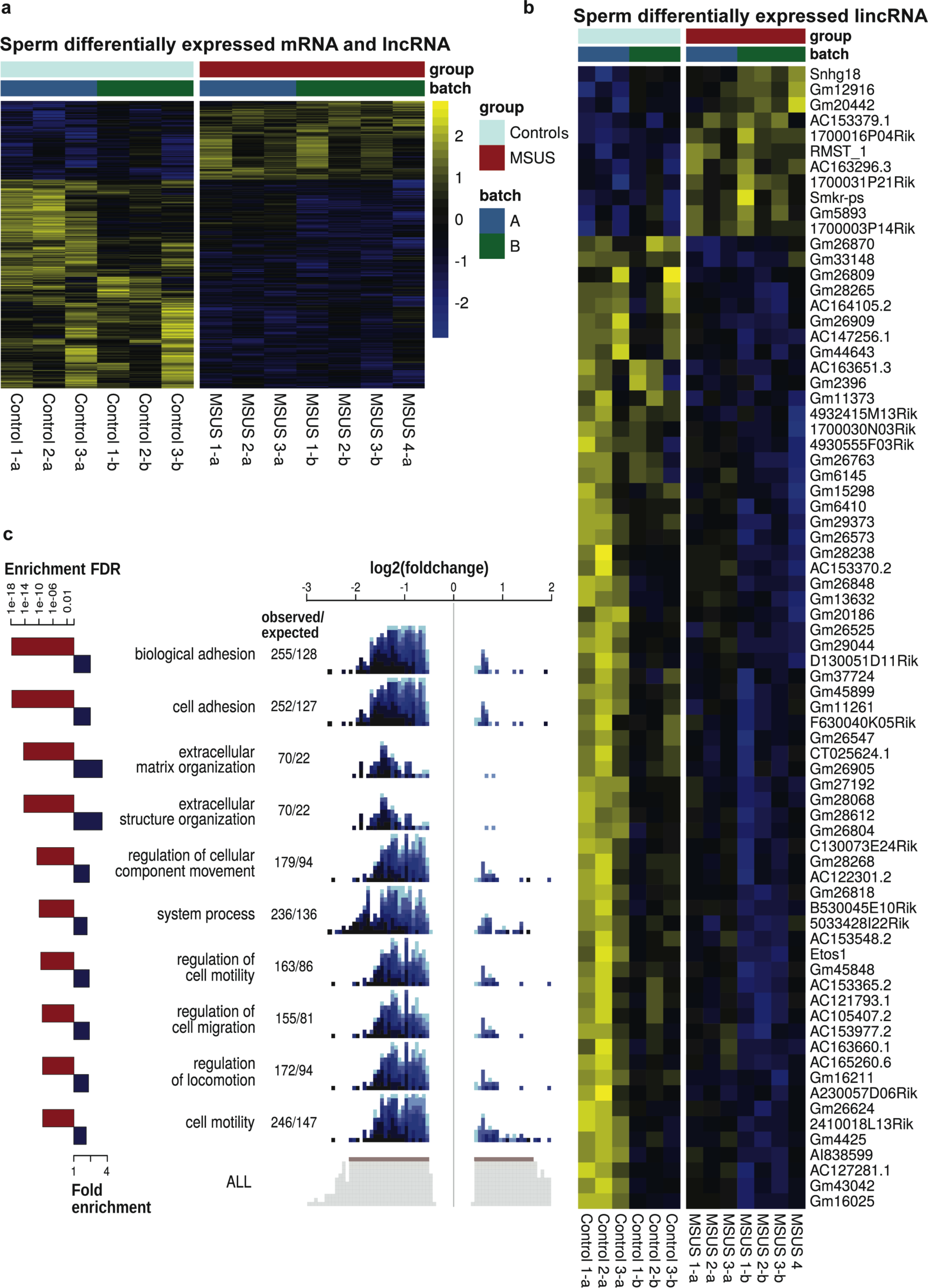
MSUS affects sperm long RNA composition. (a) Heatmap showing that MSUS induces significant changes in a range of protein coding and long non-coding RNA (lncRNA) in sperm. Plotted are the row z-scores of log normalized counts. (b) Heatmap showing single long intergenic non-coding (linc), RNA transcripts with differential accumulation in sperm of MSUS and control males (controls n=4 with 2 biological replicates, MSUS n=3 with 3 biological replicates, each replicate consists of sperm RNA pooled from 5 mice; multiple testing corrected p<0.01). (c) GO term analysis reveals enrichment of differentially expressed protein coding genes in several categories in sperm RNA from MSUS males. Red bars represent p-values of each GO term enrichment after multiple comparison correction. Blue bars represent fold enrichment of each GO term. Each blue square depicts a differentially expressed sperm transcript. Shades of blue represents the significance of its differential expression. ob= observed, exp= expected. Sequencing was done once with 2 batches of different libraries (a or b).

### The potential origin of mRNA and long non-coding RNA (lncRNA) differentially accumulated in sperm by MSUS

Based on the assumption that sperm is transcriptionally silent ^27^, we investigated the potential cellular origin of the RNA altered by MSUS in adult sperm. Meta-analyses of previously published datasets^28,29^ on the transcriptome of spermatogonia, pachytene spermatocytes and round spermatids, three different spermatogenic populations that differentiate successively during spermatogenesis to give rise to sperm revealed the presence of almost all differentially expressed gene transcripts in spermatogonia, pachytene spermatocytes (all but 3 out of 1201) and in round spermatids (all but 9 out of 1201), suggesting that transcripts differentially expressed in mature sperm originate from preceding spermatogenic cells at an earlier stage of spermatogenesis (Fig. 5). Consistent with a spermatogenic origin of some transcripts found to be altered by MSUS, analyses of Sertoli cells, somatic cells in testes, showed no expression of 112 genes, that were differentially expressed in MSUS sperm. Sertoli cells did not contain any of the 9 transcripts altered in MSUS sperm but not detectable in round spermatids, indicating potential uptake during epididymal transit (3 upregulated) or a decreased supply (6 downregulated) of immature sperm. For the 3 up-regulated transcripts, we cannot exclude uptake by for instance “plasma bridges” ^30^ after induction of expression upon exposure to MSUS in Sertoli cells.

**Figure 5.**
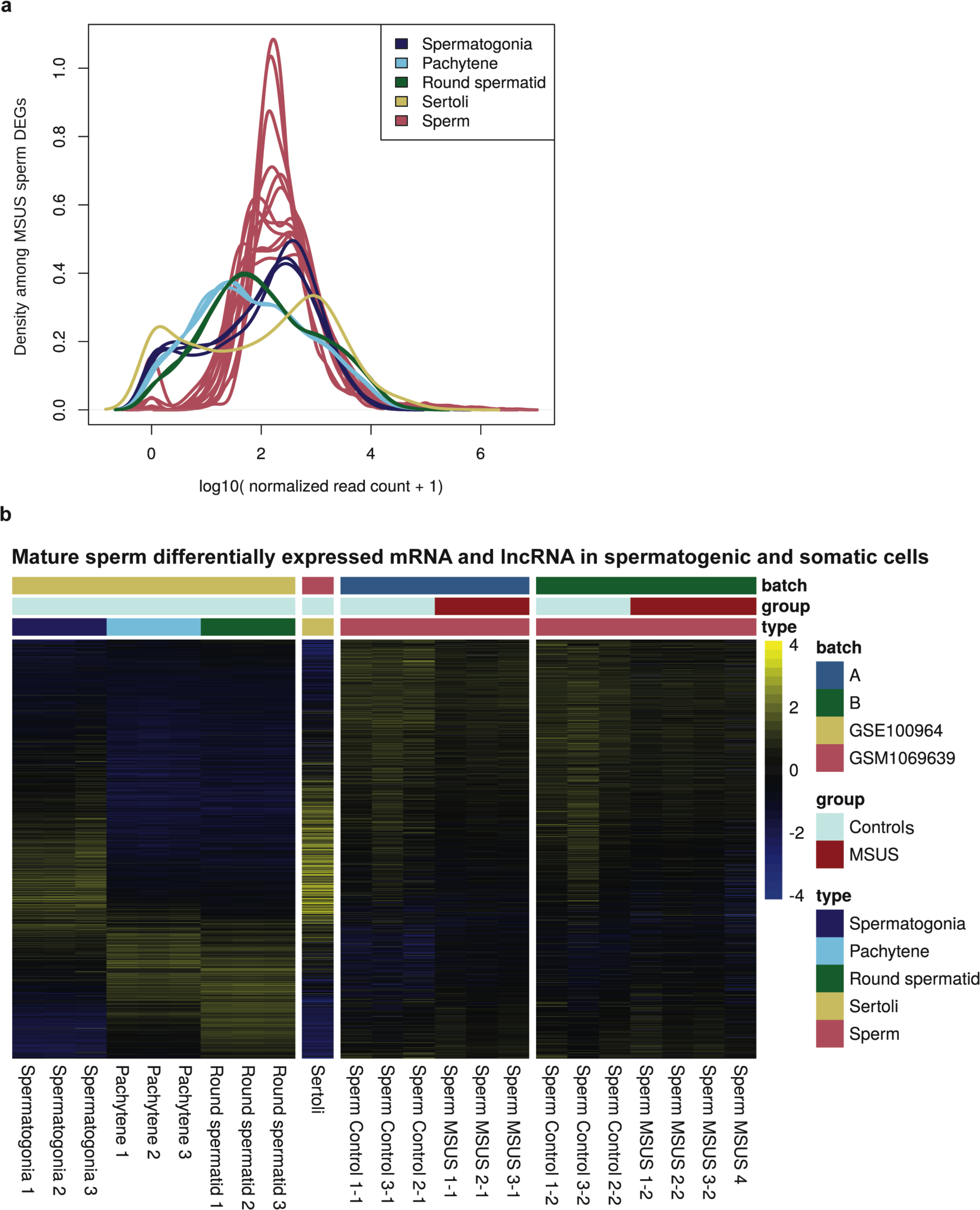
Expression of long RNA transcripts affected by MSUS in sperm at different developmental spermatogenic stages and in Sertoli cells. (a) Density plot shows the presence of mRNA and long non-coding RNA (lncRNA) found to be differentially accumulated in MSUS sperm in spermatogonia, pachytene spermatocytes (pachytene), round spermatids and Sertoli cells. (b) Heatmap shows single differentially accumulated mRNA and long non-coding RNA in F1 control and MSUS sperm in comparison to spermatogonia (3 samples), pachytene spermatocytes (3 samples), round spermatids (3 samples) and Sertoli cells (1 sample) in control conditions (published data ^28,29^). Sequencing was done once with 2 batches of different libraries (a or b). Plotted are the row z-scores of log normalized counts.

### The fate and function of mRNA and lncRNA differentially accumulated in sperm

Besides providing insight into the potential origin of altered sperm RNA, we examined the fate of these RNA after fertilization. We assessed whether RNA differentially expressed in sperm can be detected in 1-cell zygotes after mating of control females with MSUS males. Next-generation sequencing of RNA pooled from 7-10 zygotes revealed many reads mapping to protein-coding genes and long non-coding genomic regions (Supplementary Figure 4a). A majority of differentially expressed genes in MSUS sperm and zygotes had correlated fold changes in expression in zygotes (Fig. 6a), suggesting a delivery of those long sperm RNA to the zygote at fertilization. Additional differentially-expressed transcripts in MSUS zygotes, suggest early downstream effects of sperm RNA (Fig. 6b). GO term analysis of differentially expressed genes in zygotes showed an enrichment for genes involved in reproduction and import into cell among others (Supplementary Figure 10, Table S3).

**Figure 6.**
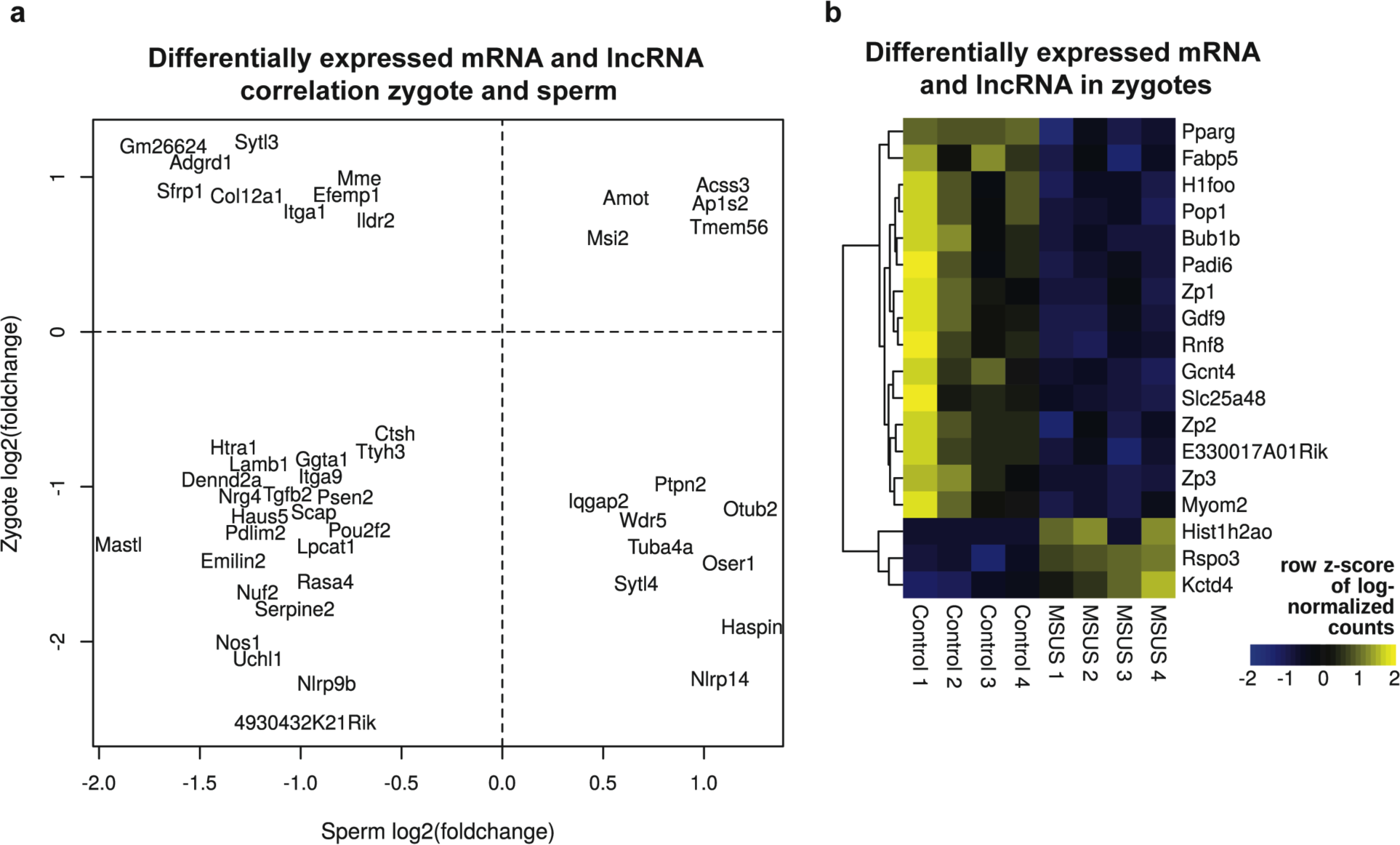
Detection of long RNA affected by MSUS in zygotes. (a) Scatter plot reveals correlation of 47 mRNA and long non-coding RNA (lncRNA) with a p-value less than 0.01 (multiple testing uncorrected) in zygotes resulting from mating of MSUS males with naïve females and a q-value less than 0.05 (multiple testing corrected) in MSUS sperm (overlap p~0.55). (b) Heatmap representing expression of 18 differentially expressed genes in F2 control and MSUS zygotes (multiple testing corrected, p<0.05). Each library reflects a pool of 7-10 zygotes (Control n=4 and MSUS n=4).

### The importance of sperm long RNA integrity to induce effects in the offspring

Because long RNA may be subjected to processing or cleavage, and generate small RNA fragments that could mediate some of the effects observed, we assessed the importance of the integrity of RNA. We fragmented long RNA from control and MSUS sperm to a size of <200nt, injected these fragmented RNA into control fertilized oocytes and assessed gene expression in resulting 4-cell embryos after injection (Supplementary Figure 4). 4-cell stage was chosen over 1-cell zygotic stage to allow enough time for injected RNA to produce their effects if any. No statistically significant alteration could be identified in embryos injected with fragmented long RNA from control or MSUS sperm (Table S4). Comparison of transcripts with putative differential expression (p>0.05 but p<0.1, not corrected for multiple comparison) in 4-cell embryos after injection of fragmented long RNA from MSUS sperm showed no changes compared to controls (Supplementary Figure 11a). Consistently, genes differentially expressed in MSUS zygotes are not dysregulated in 4-cell embryos resulting from fertilized oocytes injected with fragmented sperm long RNA (Supplementary Figure 11b). These results strongly suggest that sperm long RNA needs to be full length to induce transcriptional changes in the embryo.

## Discussion

Here we show that sperm long RNA is impacted by postnatal trauma in adulthood and provide evidence that this RNA contributes to the transmission of some of the effects of trauma in the offspring. The data also show that reproducing an excess of long RNA alone or small RNA alone by injection into fertilized oocytes is not sufficient to recapitulate all symptoms in adulthood, indicating that the combination of small and long RNA (by injecting total RNA) is necessary ^2^ and suggesting a synergistic action of long and small RNA. However, beyond a required combined action, it is also possible that a decrease (and not just an increase as reproduced by injection) in specific small and/or long RNA is also necessary to produce some of the effects, which is not recapitulated by RNA injection. In the future, manipulations mimicking down-regulation of specific small or long RNA using for instance, the CRISPR-dCas9 technology would help address this point ^31^. The results also show that MSUS effects are transmitted independently of alteration of tRNA 3’ fragments in sperm contrary to that reported in dietary models (high fat or low protein diet) ^18,23,24^. These differences in the contribution of RNA might be due to the different nature and time window of interventions, which span preconception to adulthood in the case of dietary models but only a short postnatal period (P1 to P14), 3-4 months before mating and sperm collection in our model. This highlights the complexity of the mechanisms of transmission from the directly exposed animals to the first generation of offspring. Transmission in the MSUS model, is transgenerational (up to the third generation ^2,10,12,13^). More complete phenotypic and epigenetic profiling of the dietary models would help better understand the underlying mechanisms. We speculate that a complex interplay between different sperm RNA fractions and other factors like DNA methylation, known to be altered by MSUS in adult sperm and brain across generations ^8,9,12^ may contribute to transmission, with long RNA playing an important role for some symptoms e.g. metabolic. Our data also provide evidence that a subset of altered sperm RNA is important for the early embryo since it remains altered in the zygote.

A previous study in a model of paternal low protein diet suggested regulation of MERVL elements based on data showing downregulation of MERVL targets in the offspring at embryonic stage ^23^. Our data suggest a regulation of TEs in sperm of males exposed to MSUS, potentially indicating a common denominator across different models of epigenetic inheritance. TE expression in the brain, a highly steroidogenic tissue, is responsive to acute and chronic stress ^32^. This process was suggested to involve glucocorticoid-mediated epigenetic remodeling ^33^. TE regulation in gametes, highly steroidogenic cells as well, might contribute to the observed changes in sperm in response to MSUS. Stress-induced regulation of TE expression can lead to transposition and mutagenesis in bacteria, fish and mice ^34–36^.

Long RNA affected by MSUS in sperm is expressed in round spermatids, suggesting that their alteration can occur earlier during spermatogenesis and persist until mature sperm. Further, some small RNAs are known to be transferred to maturing sperm from exosome-like vesicles called epididymosomes during epididymal transit ^37^. Such transfer may also occur in our model, and further, RNA may be provided through yet unexplored mechanisms involving “plasma-bridges” for instance ^30^. Our data further indicate that loss of RNA integrity prevents an induction of transcriptional changes in embryos associated with MSUS. The information carrier in our model is therefore different from that reported in another model of transgenerational inheritance involving c-Kit, in which truncated versions of the involved transcript could allow transmission of the phenotype ^38^. Our study hence underscores the complex repertoire of potential signals not based on DNA sequence for intergenerational transmission. Our findings might have major implications for disease susceptibility induced by early life experiences and its transmission. They may also help explore the development of RNA-based approaches to prevent the molecular transfer of the effects of early trauma onto disease risk.

## Acknowledgments

We thank Heiko Hörster for help with animal husbandry, Alexandra Sapetschnig for advice on sequencing library preparation and Tomás Di Domenico for advice on sequencing data analysis. This work was supported by the Austrian Academy of Sciences (fFORTE), the University of Zürich, the Swiss Federal Institute of Technology (grant: ETH-10 15-2), the Swiss National Science Foundation, Roche, Novartis Foundation, the Slack-Gyr Foundation, Cancer Research UK (C13474/A18583, C6946/A14492) and Wellcome (104640/Z/14/Z, 092096/Z/10/Z).

## Author contributions

K.G. carried out MSUS paradigm, metabolic measurements, sperm RNA preparation for sequencing libraries and for RNA injection into fertilized oocytes, zygote collection and sequencing library preparation. G. vS. assisted with metabolic tests and carried out MSUS paradigm with F.M.. F.M. did behavioural testing. P.L.G, K.R., W.M. and T.G. performed RNA sequencing analyses. P.P. carried out the RNA microinjection experiments. G.V. helped with sequencing library preparation. M.R. helped with sequencing results quality control. K.G., I.M.M. and E.A.M. designed the study, interpreted the results and wrote the manuscript. I.M.M. and E.A.M. raised funds to support the project.

## Additional information

Supplementary information is available.

